# Cognitive robustness in a new insect model of extended longevity

**DOI:** 10.64898/2026.01.25.701637

**Authors:** Jessica Foley, Josie McPherson, Made Roger, Fletcher J. Young, W. Owen McMillan, Stephen H. Montgomery

## Abstract

Human life expectancy has jumped by several decades in the last century, but the healthspan has not followed suit. The ageing population has come with an increased prevalence of neurodegenerative disease, and insects have proven to be valuable models of these conditions and their associated cognitive decline. The *Heliconius* butterfly genus is an emerging system in insect cognition, having recently evolved a significant expansion in neural centres of learning and memory, alongside enhanced stability of visual long-term memory in comparison with their close relatives in the Heliconiini tribe. This is linked to the cognitive demands imposed by *Heliconius*’ spatially-faithful foraging behaviour for a protein-rich diet of pollen, and co-occurs with a dramatic lifespan extension in this genus over the other Heliconiini outgroups. Here, we investigate whether the *Heliconius* cognitive healthspan is similarly discrepant, or if any cognitive decline is instead delayed in accordance with their lifespan extension. We first report evidence that investment in learning and memory circuits co-evolves with Heliconiini lifespan. We then conduct learning and memory assays across the lifespans of a representative longer-lived *Heliconius*, *H. hecale*, and the shorter-lived Heliconiini outgroup, *Dryas iulia*. Across both species, and particularly in *H. hecale*, we find evidence for cognitive robustness in late life, in contrast to evidence for widespread cognitive declines in insects. Our results add a new taxonomic order to the study of age-related memory impairment, and suggest *Heliconius* as a valuable model system for mechanistic studies of the maintenance of neurological health in the context of extended life.

## Introduction

Over the past century, improvements in public health and medical care have substantially increased human life expectancy [1]. However, this has not been matched by equivalent improvements in the healthspan, and these additional years are often spent in poor health [2]. Ageing is a prominent risk factor for many diseases [3], and with a globally ageing population [4], age-related pathologies such as cancer, cardiovascular disease, and diabetes account for the majority of global mortality [5]. In particular, the profound emotional and societal impact of neurodegenerative disease makes research into the maintenance of late-life neurological health a biomedical priority. Due to their short lifespans and tractable nervous systems, insects have long been useful models for such research, yielding seminal insights into the cognitive decline that accompanies the progression of normal ageing [6–10]. However, we lack a suitable model for understanding how nervous system function can be preserved in the context of an extended lifespan.

Butterflies in the Neotropical *Heliconius* genus form an emerging model system in the field of evolutionary cognitive ecology [11]. They have evolved an approximately 8-fold expansion of Kenyon cells [12–14], the intrinsic neurons of the mushroom body, a brain region central to learning and memory in insects [15]. Alongside this expanded cell population, the *Heliconius* mushroom body also shows evidence of specialisation towards visual processing [12]. These neural adaptations are thought to reflect the cognitive demands imposed by *Heliconius’* unique pollen-feeding behaviour, in which individuals repeatedly visit the same floral resources along stereotyped foraging routes [12, 16]. Consistent with this, *Heliconius* show a capacity to memorise spatial information [17], as well as enhanced visual long-term memory in comparison with their close relatives in the Heliconiini butterfly tribe [18]. Together with these behavioural and neural specialisations, *Heliconius* have also evolved an extended lifespan, living on average three times longer than their Heliconiini relatives [19]. These traits may be evolutionarily linked; neural expansion and longevity are often suggested to co-evolve [20–22] due to indirect selection for longer lifespan afforded by greater behavioural flexibility, a theory known as the “cognitive buffer hypothesis” [23, 24]. Empirical support for this hypothesis, however, has largely been restricted to vertebrates, with comparatively limited evidence from invertebrate systems [25].

Their extended lifespans and experimental tractability for assaying cognitive traits such as long-term memory position *Heliconius* as a promising insect model for investigating healthy neurological ageing. Given *Heliconius*’ long lives and their reliance on pollen for overall fitness [19], selection for the ability to form and retain long-term memories is expected to persist into late life. However, most animals studied to date, including many insects, show evidence for age-related memory impairments [7–10, 26–29]. We previously showed that the lengthened lifespans in *Heliconius* are accompanied by the apparent absence of physiological senescence [19], but the effect of age on cognition in these butterflies remains unknown. We therefore sought to test whether *Heliconius* exhibit a similar delay in age-related cognitive decline, conducting a series of learning and memory assays across the full lifespan of both *Heliconius hecale*, a representative longer-lived *Heliconius*, and *Dryas iulia*, a representative shorter-lived Heliconiini outgroup. Our results present the first investigation of the impact of age on cognition in Lepidoptera, and establish the *Heliconius* genus as a powerful model system for understanding the preservation of neurological health across an extended lifespan.

## Results

### Coevolution of neural investment and adult longevity

Considering that both maximum reported lifespan [19] and mushroom body size [12] vary approximately 25-fold across the Heliconiini tribe (Figure 1A), we tested for an association between longevity and mushroom body size in these butterflies [12]. Phylogenetic generalised least squares analysis (PGLS) revealed that while longevity was associated with central brain size across Heliconiini species (PGLS: *t*_22_ = 2.15, *p* = 0.043; Pearson’s *r* = 0.49), this effect was largely driven by an association between longevity and mushroom body size specifically (Figure 1B; PGLS: *t*_22_ = 3.54, *p* = 0.029; Pearson’s *r* = 0.60), with the effect of overall central brain size weakening when the mushroom body was removed from the analysis (Figure 1C; PGLS: *t*_22_ = 1.92, *p* = 0.068; Pearson’s *r* = 0.32). This provides rare evidence for such an association between brain size and longevity in invertebrates, and suggests that the enhanced cognitive abilities afforded by *Heliconius*’ mushroom body expansion [11, 12] may have facilitated the evolution of increased longevity in this genus. However, for this effect to take place, learning and memory in *Heliconius* must be robust to age-related cognitive decline.

**Figure 1:**
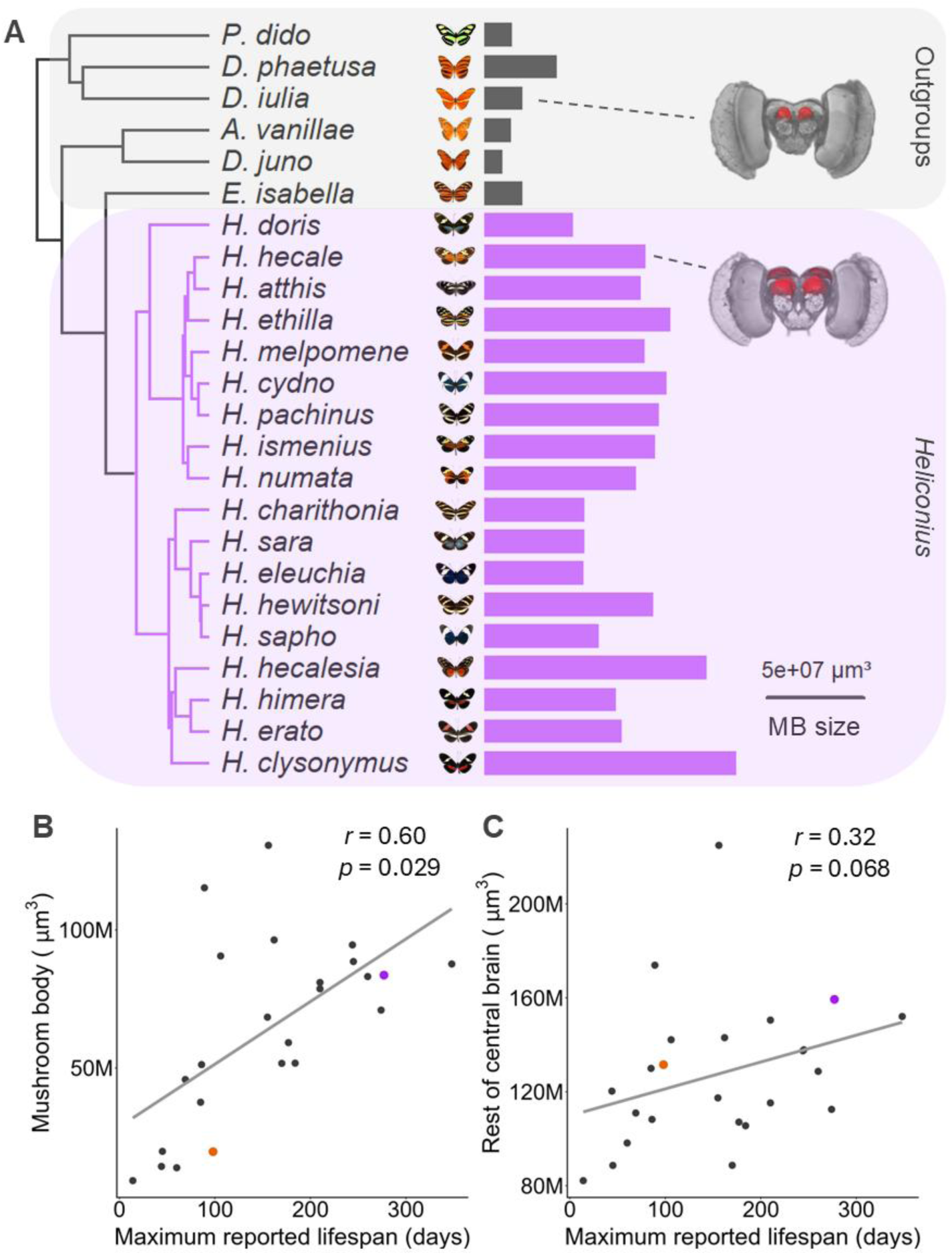
Association between mushroom body size and maximum reported lifespan across the Heliconiini butterfly tribe. **A** Phylogeny of the Heliconiini tribe showing the expansion of the mushroom body brain region in *Heliconius* in comparison with the outgroup genera. Associated bars represent mushroom body (MB) volume as reported in [12] (scale bar = 5e+07µm^3^). Sample 3D reconstructions of whole brains (anterior view) from *D. iulia* and *H. hecale* are shown for illustrative purposes with mushroom bodies highlighted in red, reproduced with permission from [12]. **B**, **C** Correlation plots between maximum reported lifespan and **B** mushroom body and **C** rest of central brain volume for all Heliconiini species shown in **A**. *p*-values are from phylogenetic generalised least squares regression with volume of the relevant brain region used as the predictor variable, and *r* is Pearson’s correlation coefficient between the two traits. Points for *D. iulia* and *H. hecale*, the two representative species used in the rest of this paper, are highlighted in orange and purple respectively for illustrative purposes. Values for maximum reported lifespan are taken from [19] and values for size of relevant brain regions are taken from [12].

### Cognitive robustness in long-lived Heliconius

To investigate if *Heliconius*’ enhanced cognition is maintained across their lengthened lifespans, and whether they showed signs of delayed cognitive senescence in comparison to their shorter-lived relatives, we repeated a previous visual long term memory test [18] at a series of ages across the lifespan of both *H. hecale*, a representative longer-lived *Heliconius*, and *D. iulia*, a representative shorter-lived outgroup.

A total of 192 individual butterflies from both species were trained for a period of 4 days to associate a particular colour of artificial feeder with either a positive (sucrose-protein solution) or negative (quinine solution) stimulus. Immediately following this training period, the strength of this trained association was tested by exposing butterflies to empty feeders of both colours and recording the proportion of feeding attempts on the “correct”, trained colour for each individual in a trained initial recall trial (“Trained”). After a period without reinforcement, butterflies were then re-tested under the same conditions to measure their long-term memory of this association over several successive trials: first after an 8-day wait (“LTM1”); then after a further 4-day wait (“LTM2”), and finally after another 4-day wait (“LTM3”). To test for any evidence of an age- related memory impairment, butterflies were entered into the training and testing protocol at a range of ages designed to cover the natural lifespan *D. iulia* (completing at up to 54 days old, almost 100% of their captive lifespan [Figure S1]) and the extended lifespan of *H. hecale* (completing at up to 90 days old, approximately 84% of their captive lifespan [Figure S1]) (Figure 2A).

**Figure 2:**
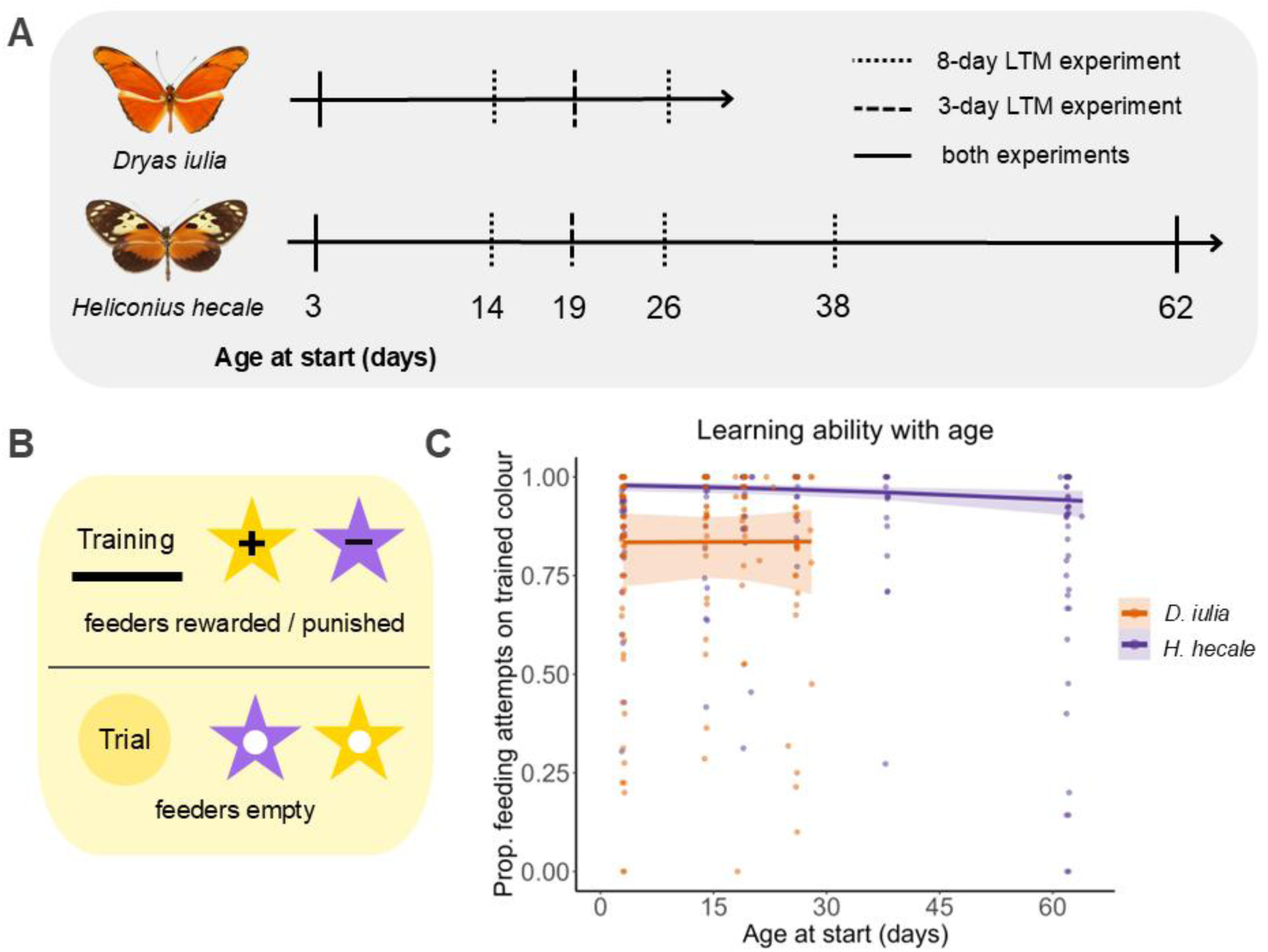
Impact of age on performance in a visual associative learning assay in *H. hecale* and *D. iulia*. **A** Age points at which individuals were entered into the testing protocol for each species. For the 8-day long-term memory (LTM) assay, butterflies were entered into the testing protocol at 3, 14, and 26 days (for both *D. iulia* and *H. hecale*), and also at 38 and 62 days (for the longer-lived *H. hecale*). For the 3-day LTM assay, butterflies were entered into the testing protocol at 3 and 19 days (for both *D. iulia* and *H. hecale*), and also at 62 days (*H. hecale*). Learning ability shown in **C** was assessed using a pooled dataset across all age groups as the 8-day and 3-day protocols up to this point (“Trained”, the initial recall trial) did not differ between the two assays. **B** During training periods, butterflies were presented with a rack of purple and yellow artificial feeders, with one colour filled with a sucrose-protein solution (“+”, the positive stimulus) one colour filled with a quinine solution (“–“, the negative stimulus). The assignation of a colour to a positive or negative stimulus varied depending on the result of each individual’s naïve preference test (see Methods; Figure 3A; Figure 3D), with individuals trained to associate the positive stimulus with the opposite colour to their innate preference. During trial periods, butterflies were presented with a rack of empty purple and yellow artificial feeders to assess the strength of the learned association. **C** Performance in the trained initial recall trial (used as a metric of learning ability) across the lifespan for *H. hecale* and *D. iulia*. “Age at start” is based on the age of entry to the protocol, with learning ability assessed 4 days later (Figures 3A & 3D, “Trained”). Dots represent individual data points indicating the proportion of feeding attempts made on the trained colour during the trained initial recall trial. Regression lines, with 95% confidence intervals, are from GLMM analysis and show predicted learning ability with age for each species.

Combining all data on performance in the initial recall trial across two experimental cohorts, our experiments confirmed the improved visual learning of *H. hecale* over *D. iulia* [18], choosing the trained colour with a predicted accuracy of 96.02% as compared with 87.49% in the trained initial recall assay (Figure 2C; *z* = 4.82, *p* < 0.001). They also showed no evidence for an impact of age on learning ability in *D. iulia* (Figure 2C; χ^2^ < 0.001, *p* = 0.978), but revealed slight impairments in visual learning with age in *H. hecale*, with butterflies at the oldest age groups showing a 5.52% decrease in accuracy compared to the younger groups in choosing the trained colour (Figure 2C; χ^2^ = 8.05, *p* = 0.005). However, this impact of age appears to only manifest at the oldest ages measured in *H. hecale*, disappearing in an inter-specific model when only the younger age groups (those within the natural lifespan of *D. iulia*) are retained for comparison with *D. iulia* (χ^2^ = 0.17, *p* = 0.683).

Despite this slight impairment in learning, we found no evidence for a main effect of age on long-term memory after 8 days in *H. hecale* (Figure 3B; χ^2^ = 1.43, *p* = 0.231), although there was a significant interaction between age and sex (χ^2^ = 4.15, *p* = 0.042), where older males showed a decline in memory performance with age (reduction in accuracy of 10.28% over the full time course), whereas older females showed an improvement (increase in accuracy of 12.56%). This may reflect an artefact of the experimental environment – for example, the lack of host-plants provided to experimental females may have led to oocyte resorption and redistribution of resources, as has been shown in some *Heliconius* species [30], potentially protecting against age-related declines. Regardless, the lack of a main effect of age in individuals tested up to 90 days old suggested a slowed cognitive senescence in *Heliconius* in association with their lengthened lifespans.

**Figure 3:**
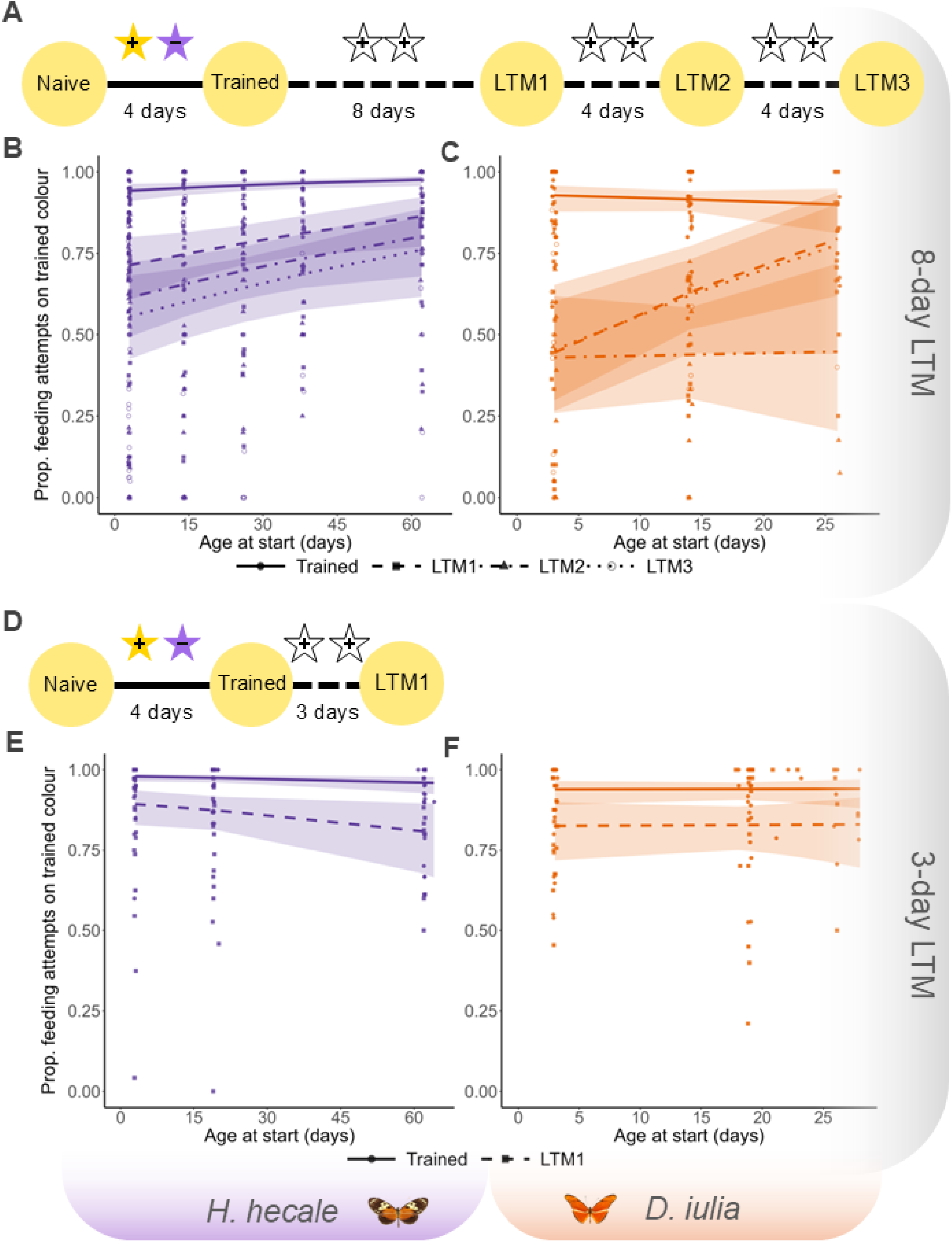
Cognitive robustness in ageing *H. hecale* and *D. iulia*. **A**, **D** Testing protocol for the 8-day and 3-day long-term memory (LTM) assay, respectively. Yellow circles denote trials, during which butterflies were presented with a rack of empty purple and yellow artificial feeders and the proportion of feeding attempts on each colour was noted. Solid lines denote training periods, during which butterflies had access to purple and yellow artificial feeders containing either a sucrose-protein solution (“+”) or a quinine solution (“–“). Dashed lines denote wait periods, during which butterflies were provided with white artificial feeders filled with a sucrose-protein solution (“+”). “Naive” = naïve preference trial, “Trained” = trained initial recall trial, “LTM” – long-term memory (suffixes 1-3 denote successive trials). **B**, **C** Performance at each trial in the 8-day LTM assay across the lifespan for **B** *H. hecale* and **C** *D. iulia*. “Age at start” is based on the age of entry to the protocol, with the “Trained” age 4 days later than the age at start, the “LTM1” age between 14 and 20 days later than the age at start, the “LTM2” age between 18 and 24 days later than the age at start, and the “LTM3” age between 22 and 28 days later than the age at start. **E**, **F** Performance at each trial in the 3-day LTM assay for **E** *H. hecale* and **F** *D. iulia*. The “Trained” age is 4 days later than the age at start, and the “LTM1” age is between 9 and 15 days later than the age at start. Points represent individual data points indicating the proportion of feeding attempts made on the trained colour during each of the four trials. Regression lines, with 95% confidence intervals, are from GLMM analysis and show predicted performance on each trial. Trial is delineated by line type and by point shape to aid visual distinction.

### A potential facultative role of cognitive robustness in the evolution of longevity

We next sought to assess whether this robustness was an evolved trait in *Heliconius* by comparing the impact of age on long-term memory in the shorter-lived *D. iulia*. However, after the initial 8-day period without reinforcement, *D. iulia* had largely forgotten the learned association, not deviating significantly from random choice in the first long-term memory trial (Figure 3C; *z* = 0.10, *p* = 0.918). This rendered comparisons with older age groups, and with *H. hecale*, uninformative. In these models, there was a significant interaction between age and trial (χ^2^ = 10.37, p = 0.016), because older *D. iulia* butterflies performed better than their younger counterparts on LTM1 (Figure 3C; z = 2.97, p = 0.003); however, this age effect was not found in any other trial (although a similar, nonsignificant trend was observed at LTM3; Figure 3C).

The comparison with *Heliconius*’ shorter-lived relatives was crucial for our understanding of the evolutionary trajectory of this apparent cognitive robustness, and so we conducted a new learning and memory experiment with a reduced wait period and a simplified structure (Figure 3). A total of 137 butterflies of both species were trained under the same conditions as the previous experiment, but were only subjected to a single long-term memory trial (“LTM1”) after a wait period of just 3 days as opposed to 8 days (Figure 3D), at which point pilot experiments suggested both species retained the learned colour association. Once again, data from these experiments showed no evidence for an age-related memory impairment in *H. hecale*, with 77-day old butterflies performing just as well as their younger 18- and 34-day old counterparts in the 3-day long-term memory trial (Figure 3E; χ^2^ = 2.49, *p* = 0.115). Considering the evidence for delayed physiological senescence in *H. hecale* in comparison with the declines seen in aged *D. iulia* [19], we had expected to find similar patterns in cognitive senescence between these two species. Interestingly, however, in the shorter-lived *D. iulia*, we also found no effect of age on 3-day long-term memory, with no difference in performance between 18- and 34-day old butterflies (Figure 3F; χ^2^ < 0.01, *p* = 0.957).

The apparent cognitive robustness demonstrated by these two insect species is particularly surprising considering the evidence for widespread age-related learning and memory impairments in other insects (Table S1; [7–10, 27–29]). This may reflect modality-specific impacts of age, with most evidence for age-related impairments of learning and memory in insects coming from olfactory conditioning paradigms [7, 8, 10, 27–29] as opposed to visual memory as is investigated here. However, this seems unlikely given the multi-modal basis of most insect behaviours [31]. Age-related cognitive decline is also not found in all insects; long-lived ant workers show no evidence of neurological or behavioural senescence [32], and the evidence for age-related memory impairments in honey bee workers is equivocal, with most impairments appearing to be related more to social role and degree of foraging rather than chronological age [33–36]. Nevertheless, the cognitive robustness demonstrated by aged *D. iulia* and by age- matched *H. hecale* may have implications for the evolution of long life in *Heliconius*. In particular, these results suggest that the last common ancestor of both *H. hecale* and *D. iulia*, likely a short-lived non-pollen-feeder, was similarly cognitively robust. This may have facilitated lifespan extension in *Heliconius*, with the adaptive benefit of longevity uninhibited by declining fitness benefits in longer-lived individuals due to cognitive deficits.

### Selective disappearance does not explain performance of old-age groups

We considered it possible that this apparent cognitive robustness was instead due to compositional differences between age classes as a result of poorer-performing individuals dying during the experiments, a phenomenon known as selective disappearance [37]. However, the typical approaches used to correct for this were not available to us, requiring either repeated measures of the same trait across multiple ages for each individual [38, 39], or unbiased estimates of individual longevity [40]. While we did collect survival data for all individuals in our dataset (see Supplementary Results), much of it was right-censored after the point at which individuals completed the learning and memory protocol (see Methods), meaning longevity estimates are incomplete. If selective disappearance was sufficient to completely mask an impact of age (as we have largely found), there would need to be a very low probability of survival to our oldest age groups, and an extremely tight correlation between survival and cognitive performance. We therefore rationalised that if cognitively “worse” individuals were indeed dying earlier, we should be able to see this effect using traditional survival analysis. To test this, we created Cox proportional hazards models to assess whether cognitive performance predicted survival in either species. To avoid any bias introduced by selective disappearance, we conducted this analysis only on the youngest age group of each species, using performance in the trained initial recall trial as a proxy for cognitive ability. Our results showed that cognitively “worse” individuals were in fact less likely to survive in *D. iulia* (*z* = −2.48, *p* = 0.013), with an individual scoring 50% in the trained initial recall trial predicted to have approximately double the mortality risk in comparison with an individual scoring 100% (hazard ratio_−0.5_ = 2.25). Therefore in *D. iulia*, it is possible that the effects of selective disappearance may have been masking a cognitive decline in older individuals, although we are unable to quantify the extent of this with the data currently available to us. Intriguingly, however, we found no evidence for a similar effect in *H. hecale*. Performance in the trained initial recall trial was not a significant predictor of survival in this species, with cognitively “worse” individuals no more likely to die, and in fact, the directionality of the trend was reversed (hazard ratio_−0.5_ = 0.27, *z* = 1.69, *p* = 0.091). This further supports our conclusion that, in contrast with most other insects studied, *H. hecale* individuals maintain cognitive performance even at the oldest ages tested, far beyond the median lifespans of their closest relatives.

## Discussion

Our results present unusual evidence for the maintenance of cognition in late life in two species of tropical butterfly. The evidence for cognitive robustness in *Heliconius* is particularly interesting considering the high energetic cost of neural tissue [41] and the cognitive processes it mediates [42–44]. While there is support for the co-evolution of brain size and lifespan in many vertebrate taxa [20–22], the compromised energy budget of relatively large-brained species [25] might instead be expected to lead to trade-offs with lifespan [45], with fewer energetic resources available to allocate towards longevity and somatic maintenance [46]; a pattern which has indeed been shown in several studies on fish [47–49]. Such trade-offs would theoretically only magnify with the elongation of lifespan, as has been demonstrated in *Drosophila melanogaster*, where artificial selection for longer lifespans comes at the cost of reduced olfactory learning performance [50]. Any potential cognitive robustness in *Heliconius*’ shorter-lived ancestors may therefore have still faltered when the limited resources for such maintenance were stretched out over longer lives. However, the evolution of a unique pollen-feeding behaviour in *Heliconius* appears to have loosened these constraints [19], with pollen-derived lipids and amino acids, both important energy storage compounds in insects [51, 52], allowing for an increased energy budget in these butterflies. This may have allowed for greater investment in neural tissue, as is seen in the *Heliconius* mushroom body expansion [12], but also in the lifetime maintenance of the cognitive abilities this expansion facilitates, as seen in the cognitive robustness of ageing *H. hecale*.

Our results build upon previous work showing improved visual long-term memory in *Heliconius* by demonstrating a retention of a visual association in *H. hecale* up to 18 days after first learning it. This exceeds the previous record of 13 days in *Heliconius melpomene* [18], and is to our knowledge, the longest visual memory recall found in an insect – although longer olfactory and spatial memories have been reported in crickets [53], honey bees [54], and red wood ants [55, 56]. The superior visual long-term memory in *Heliconius* is demonstrated following an 8-day wait period without reinforcement, at which point non-pollen-feeding Heliconiini outgroups no longer retain the learned association [18]. Our results support this finding (Figure 4A), but show that this cognitive advantage in *Heliconius* emerges only over longer timeframes, with *H. hecale* performing no better than *D. iulia* on the long-term memory trial when the wait period is reduced to only 3 days (Figure 4B).

**Figure 4:**
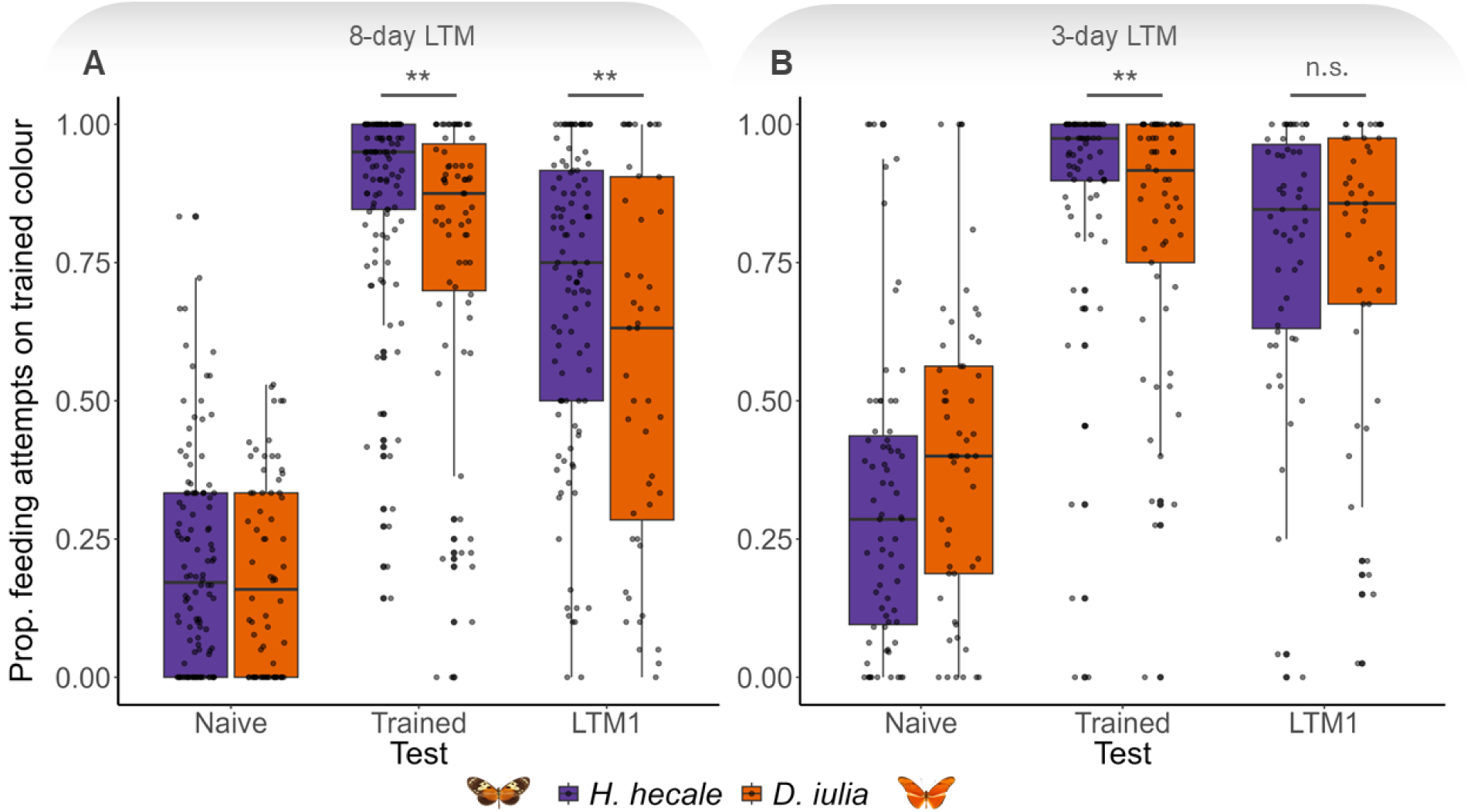
Performance of *H. hecale* and *D. iulia* in 8-day and 3-day long-term memory assays. Performance of both *H. hecale* and *D. iulia* (full datasets, pooled across all age groups) in long-term memory assays following (a) an 8-day wait period and (b) a 3-day wait period. To aid comparison between the two protocols, only the first long-term memory test (LTM1) from the 8-day protocol is presented. Dots represent individual data points indicating the proportion of feeding attempts made on the trained colour during each of the trials. “Naive” = naïve preference trial, “Trained” = initial post-training recall test, “LTM1” – long-term memory trial. n.s.: p > 0.05; *: p < 0.05; **: p < 0.01; ***: p < 0.001.

At the physiological level, this difference might reflect different phases of long-term memory, such as those characterised in detail in *Drosophila* [57]. In fruit flies, olfactory classical conditioning results in several types of consolidated memory [58], one of which is dependent upon protein synthesis, and persists for at least 7 days (protein synthesis-dependent long-term memory, or PSD-LTM), and one of which does not require protein synthesis, and largely decays within 3 days (anaesthesia-resistant memory, or ARM) [59]. Though deeper pharmacological dissection (i.e. to discern anaesthesia sensitivity and protein synthesis requirements) would be required to accurately place our results into this context, the similar timescales between our 8-and 3-day long-term memory experiments and the retention periods in *Drosophila* PSD-LTM and ARM present an interesting case for such a distinction between memory phases in Heliconiini butterflies. Specifically, the requirement for protein synthesis in PSD-LTM might suggest a physiological basis for our finding of improved 8-day, but not 3-day, long-term memory in *H. hecale*. Learning and memory are costly [42, 44], and PSD-LTM is more stable than ARM but also more expensive, incurring higher fitness costs [43]. This expenditure may be more achievable for *Heliconius* than for other Heliconiini, however, due to the dietary amino acids provided by their adult pollen-feeding behaviour [16], which may contribute to increased investment in PSD-LTM in these butterflies. As *Heliconius* may repeatedly return to known pollen sources for a period of months [60], the superior performance after an 8-day (and even 18-day) wait in *H. hecale* over *D. iulia* likely reflects an increased investment into an ecologically-relevant behaviour. Such persistent memories would indeed be particularly important in *Heliconius* considering their lengthy lifespans. For the shorter-lived, non-pollen-feeding Heliconiini outgroups, which are suggested to have less stable home ranges [61], long-term memory after such periods might be an unnecessarily expensive investment.

This work represents the first investigation of the impact of age on cognition in Lepidoptera, offering new insights into a novel insect system emerging as a valuable model in evolutionary cognitive ecology [11] and the comparative biology of ageing [19]. Our finding of an association between lifespan and brain size in the Heliconiini tribe provides rare evidence for such an association outside of vertebrates [25], and lends support to the cognitive buffer hypothesis by linking lifespan to a specific neural expansion previously associated with enhanced cognition [18]. The capacity of *H. hecale* to perform cognitively demanding tasks, such as the consolidation of new long-term memories, at ages far beyond the lifespan of their close relatives, presents *Heliconius* as a powerful model for the maintenance of neural processes in the context of an extended lifespan. In an ageing global population, the preservation of cognitive faculties late in life has become a biomedical and economic imperative [62, 63]. Future research on the mechanistic basis of extended neuronal longevity in *Heliconius* could provide a much-needed window into the biology of healthy neurological ageing.

## Acknowledgements

We are very grateful to the Ministerio del Ambiente in Panama for permission to collect butterflies used in these experiments, and to the Smithsonian Tropical Research Institute for providing the facilities to carry out much of this research. We would like to thank Laura Hebberecht-Lopez, Lina Melo, Cruz Batista, and Oscar Paneso for their assistance with rearing at the insectaries. This work was supported by a Smithsonian Tropical Research Institute Short Term Fellowship and a DTP Studentship as part of the GW4 BioMed MRC DTP (MR/N0137941/1) awarded to the Universities of Bath, Bristol, Cardiff and Exeter from the MRC/UKRI (J. F.), and a NERC Independent Research Fellowship (NE/N014936/1) and ERC Starter Grant (758508) (S.H.M.), and funding from the US National Science Foundation (IOS 2110532) and the Smithsonian Tropical Research Institute (W.O.M.).

## Author contributions

Conceptualization: S.H.M. Methodology: J.F., F.J.Y., and S.H.M. Investigation: J.F., J.M., M.R., and F.J.Y. Formal analysis: J.F. Resources: W.O.M. and S.H.M. Visualization: J.F. Supervision: W.O.M. and S.H.M. Funding acquisition: W.O.M. and S.H.M. Writing – original draft: J.F. and S.H.M. Writing – review and editing: All authors.

## Methods

### Butterfly husbandry and survival data collection

All experimental individuals were reared from stock populations housed in outdoor insectaries at the Smithsonian Tropical Research Institute, Gamboa, Panama. Stock populations of *Heliconius hecale melicerta* and *Dryas iulia* were established from wild-caught individuals captured within a 2km radius of Gamboa during two separate field seasons, running from March-June 2019 and January-November 2022, under permit number SE/A-82-19. Stocks were maintained in 2m (L) x 2m (W) x 2m (H) cages containing artificial feeders filled with a 20% w/v sucrose and 10% w/v organic, pesticide-free bee pollen solution, changed every 2 days. All stock cages contained *Passiflora* host-plants as well as flowering *Palicourea, Lantana,* and *Stachytarpheta* plants, and *H. hecale* stocks were also provided with fresh *Psiguria* flowers daily. Eggs were collected daily and larvae were reared in pop-up cages. feeding *ad libitum* on shoots of their preferred *Passiflora* host-plants: *P. biflora*, *P. auriculata, P. pittieri,* or *P. edulis* for *D. iulia*; and *P. nitida, P. riparia,* or *P. vitifolia* for *H. hecale*.

Upon eclosion, individuals were sexed, marked with a unique ID, and placed in a pre-training experimental cage, which had white feeders containing 0.5ml of a sucrose-protein solution (25% w/v sucrose, 5% w/v Vetark Critical Care Formula). Butterflies were taught to feed on these artificial feeders by manually extending their proboscis into the solution. Individuals remained in these cages until beginning the testing protocol. All experimental cages (pre-training, training, testing, and wait cages) measured 2m (L) x 3m (W) x 2m (H) and contained a single rack of 24 star-shaped feeders arranged in a 4 x 6 grid. All solutions in the experimental cages were replaced daily. All experimental cages also contained a single non-flowering *Palicourea elata* plant provided as a roosting site for butterflies.

Cages were checked daily for dead individuals and ages at death were noted for survival analysis. After completing the testing protocol, most butterflies from the 2019 field season were dissected for further neuroanatomical analysis. Butterflies which had not yet completed the protocol by the end of the 2019 field season were released into the wild. Butterflies completing the testing protocol from the 2022 field season were left to live out their natural lives in stock cages in the Gamboa insectaries, and were not always monitored for deaths. Butterflies that had been predated upon, dissected, released, or moved to an unmonitored cage were noted for censorship in survival analysis.

### Learning and memory assays

For all learning and memory assays, butterflies were trained to associate either a purple or yellow colour with a positive or negative stimulus, following the protocol established by Young *et al*. (2024) [18]. Upon entering the testing protocol, butterflies were first subjected to a naïve preference trial (“Naive”) to identify any innate preference for either colour. If an individual demonstrated a preference for one colour over another, they were trained to associate the opposite colour with a positive stimulus (the trained colour) and their preferred colour with a negative stimulus (the untrained colour). If an individual demonstrated no preference for either colour, the trained colour was assigned randomly using a coin-toss.

Training cages contained an equal distribution of both purple and yellow artificial feeders, arranged at random on the 4 x 6 rack. Feeders of the trained colour were filled with 0.5ml of the sucrose-protein solution (positive stimulus) and feeders of the untrained colour were filled with 0.5ml of a saturated quinine solution (negative stimulus). Butterflies were kept in the training cage for 4 days, allowing them time to develop a learned association between the trained colour and the sucrose-protein reward. Following the 4-day training period, butterflies were moved to a test cage and subjected to an initial recall trial (“Trained”) to assess how well they learned the association (Figure 2B). Upon completion of this trial, individuals were moved to a holding cage (the “wait” cage) containing only white feeders filled with the sucrose-protein solution, until beginning the long-term memory trials as described later in this section.

All testing (naïve preference trials, trained initial recall trials, and long-term memory trials) was conducted for a 4-hour interval between 08:00 and 12:00. During testing, butterflies were presented with an equal distribution of purple and yellow feeders, arranged at random on the 4 x 6 rack. The feeders in the testing trials were left empty to ensure that feeding attempts were made based only on visual cues. The butterflies were filmed for the 4-hour interval using a GoPro Hero 6 camera mounted on a tripod, and videos were manually reviewed, recording the number of attempts made by each individual on each colour. At the end of the testing interval, individuals were returned to the cage they had been housed in just prior to the testing period: the pre-training cages for the naïve preference trial, the training cages for the trained initial recall trial, or the wait cages for the long-term memory trials. If a butterfly had made at least 15 feeding attempts, it was considered to have completed testing, and was moved on to the next stage of the protocol. If a butterfly made fewer than 15 feeding attempts, it was tested again the following day, and again for up to a maximum of 4 days, until it had made a cumulative 15 feeding attempts. If it had not made at least 15 attempts by the end of the 4^th^ day of training, it was moved on to the next stage of the protocol regardless. In total, data on performance in the trained initial recall trial (analysed as a metric of learning ability) were collected for 195 individuals of *H. hecale* and 132 individuals of *D. iulia*. A full breakdown of sample size per species and age group may be found in Tables S2 and S3.

### 8-day long-term memory assay

Butterflies from the 2019 field season were assessed for long-term memory of the colour association after an 8-day wait period, chosen for its ecological relevance (see Results and Discussion), and based on pilot data from another lab member showing superior 8-day long-term memory in *Heliconius* when compared with other Heliconiini (ultimately published in [18]). Following completion of the trained initial recall trial, butterflies were moved to wait cages containing only white feeders filled with the sucrose-protein solution, and left there for a period of 8 days. Following this 8-day wait period, during which the initial trained colour association was not reinforced, butterflies were tested for retention of this association in the first long-term memory trial (“LTM1”), following the testing procedure described above. Upon completing LTM1, butterflies were placed back into the wait cages for a further 4 days, before then being tested once again for their long-term memory of this association (“LTM2”). Upon completion of LTM2, butterflies were once more moved to the wait cages for a further 4 days, before then being tested one final time for retention of the colour association (“LTM3”) (Figure 3A).

To test for an impact of age on learning and long-term memory, butterflies were entered into this protocol at different ages (measured in days post-eclosion), chosen to span the insectary adult lifespan of both *D. iulia* and *H. hecale* [19]. *D. iulia* butterflies, being shorter-lived, were entered into the protocol at three different age points, starting the protocol at 3, 14, and 26 days old (Figure 2A). Longer-lived *H. hecale* individuals were entered into the protocol at five different age points, the first three corresponding to the age points in *D. iulia*, plus an additional two beyond these, starting the protocol at 3, 14, 26, 38, and 62 days old (Figure 2A). Depending on how long butterflies took to reach 15 attempts in the testing periods (between 1 and 4 days), individuals were between 14 and 20 days older than these initial age points by the time they were tested at LTM1, between 18 and 24 days older by the time they were tested at LTM2, and between 22 and 28 days older by the time they were tested at LTM3 (Figure 3A). In total, data on 8-day long-term memory performance were collected for 121 individuals of *H. hecale* and 71 individuals of *D. iulia*. A full breakdown of sample size per species, trial, and age group may be found in Table S2.

### 3-day long-term memory assay

Results from the 8-day long-term memory assay conducted in 2019 proved difficult to assess for any differential impact of age between *H. hecale* and *D. iulia*, because after an 8-day wait *D. iulia* individuals had largely forgotten the learned association (see Results and Discussion). Therefore the assay was repeated in a second field season in 2022 with a reduced wait period of 3 days before the long-term memory trial, chosen because pilot experiments showed that by this point, butterflies of both species still retained the learned colour association. The structure of the testing protocol was also simplified, eliminating the second and third long-term memory trial, retaining only the first (“LTM1”); and reducing the age groups to just two age points in *D. iulia* (entering the protocol at 3 and 19 days old) and 3 age points in *H. hecale* (entering the protocol at 3, 19, and 62 days) (Figure 3D). The chosen age points were initially intended to correspond to those of the 8-day experiment in 2019, but overall survival was lower in this cohort (see Supplementary Results), and the 26-day age point proved too challenging to reach in *D. iulia*. Therefore the oldest age point in *D. iulia*, and its corresponding age point in *H. hecale*, were reduced by a week to 19 days, though data from 8 *D. iulia* individuals that successfully completed the protocol having begun it at 26 days were also included in analysis. Depending on how long butterflies took to reach 15 attempts in the testing periods (between 1 and 4 days), individuals were between 9 and 15 days older than these initial age points by the time they were tested at LTM1 (Figure 3D). In total, data on 3-day long-term memory performance were collected for 76 individuals of *H. hecale* and 61 individuals of *D. iulia*. A full breakdown of sample size per species, trial, and age group may be found in Table S3.

### Statistical analyses

All statistical analyses were conducted using R v.4.3.1 [64].

#### Association between brain size and longevity

Associations between Heliconiini maximum reported lifespan and size of relevant brain regions were tested using phylogenetic generalised least squares (PGLS) regression, using the gls function in the R package *nlme* v3.1-168 [65], with a Brownian-motion correlation structure implemented via *ape* v5.8-1 [66]. In line with the cognitive buffer hypothesis, which predicts that increased neural investment should lead to selection for longer life, brain region volume was used as the predictor variable in all models. Descriptive statistics for the effect size of the correlation were estimated using Pearson’s correlation coefficient. Volumetric data on Heliconiini brain size (measured in µm^3^) were taken from [12], subtracting mushroom body volume from the total central brain volume to obtain values for “Rest of central brain” in Figure 1. Values for Heliconiini maximum reported lifespan (measured in days) were taken from [19]. Data on both brain size and maximum lifespan were available and analysed for 24 Heliconiini species: *Agraulis vanillae*, *Dione juno*, *Dryadula phaetusa*, *Dryas iulia*, *Eueides isabella*, *Heliconius atthis*, *Heliconius charithonia*, *Heliconius clysonymus*, *Heliconius cydno*, *Heliconius doris*, *Heliconius eleuchia*, *Heliconius erato*, *Heliconius ethilla*, *Heliconius hecale*, *Heliconius hecalesia*, *Heliconius hewitsoni*, *Heliconius himera*, *Heliconius ismenius*, *Heliconius melpomene*, *Heliconius numata*, *Heliconius pachinus*, *Heliconius sapho*, *Heliconius sara,* and *Philaethria dido*.

#### Survival analyses

Kaplan-Meier survival curves and corresponding median survival estimates were generated using the R package *survival* v3.5-5 [67] for both the 2019 and 2022 cohorts in order to justify the choice of age groups (Supplementary Results). To investigate the possibility of selective disappearance, Cox proportional hazards models were created using the R package *coxme* v2.2-18 [68]. These were run on the youngest age group of each species, including the proportion of correct feeding attempts (as a numerical value bounded between 0 and 1) for each individual in the trained initial recall trial as a predictor of survival.

#### Cognitive senescence

For each of the cognitive traits assessed (learning ability, 8-day long-term memory, and 3-day long-term memory), three models were created: two single-species models, fit to datasets containing data from all age groups for each of *H. hecale* and *D. iulia*; and one inter-specific model, fit to a dataset containing data from both species, up to the maximum age for which there was data in *D. iulia*. Age was treated as a continuous variable in all models, and was initially recorded in days, but converted to weeks during analysis to aid model convergence. For all models, stepwise backward model selection was implemented by performing an analysis of variance (ANOVA) on the model with and without a named predictor, to see if inclusion of the predictor significantly improved model fit. The final “best” model was then compared with an intercept-only “null” model to confirm its validity. An ANOVA was then run on the final best model to report significance of named predictors. Diagnostics for all models were conducted using the using the R package *DHARMa* v0.4.6 [69]. In cases where overdispersion was detected, this was corrected for using an observation-level random effect [70]. Model predictions for visualisation of results were created with the aid of the R package *ggeffects* v1.3.0 [71].

Learning ability was analysed using generalised linear mixed models created in the R package *lme4* v1.1-34 [72], including an observation-level random effect due to overdispersion in the original general linear models. These were fit to a binomial distribution, which treated feeding attempts on the “correct”, trained colour as successes, and feeding attempts on the “incorrect”, untrained colour as failures. Learning ability was assessed on a pooled dataset containing individuals from both the 2019 and 2022 field seasons in order to maximise statistical power, as the protocols up to this point (“Trained”, the initial recall trial) did not differ between the two cohorts (Figure 3A, Figure 3D).

For assessment of long-term memory performance, any individuals scoring less than 50% in the trained initial recall trial were removed from analysis, as they had not learned the colour association, and therefore their memory of this association could not be meaningfully assessed. Long-term memory performance was also analysed using generalised linear mixed models in *lme4*, this time including individual ID as a random effect to account for repeated measurements of individuals across trials. These were also fit using a binomial distribution, again treating the number of correct versus incorrect colour choices as the response variable. For all learning and memory assays, up to 40 feeding attempts in each individual on each trial were included in analysis, as described in [18]. Though it is included in some plots (Figure 4) to aid visualisation, performance in the naïve preference trial was not used for analysis. This is because while it is measured on the same response variable, it measures a very different biological process, capturing the strength of an innate colour preference as opposed to the strength of a learned association.

For the learning assays, all single-species “full” models included age, sex, and their interaction as candidate fixed effects. For the long-term memory assays, all single-species “full” models included age, sex, trial (i.e. “Trained”, “LTM1”, etc.), and the 2-way interactions between each of these as candidate fixed effects. All inter-specific “full” models then tested for interactions of these candidate fixed effects with species. If an effect seen in a single-species model is described as “disappearing” in an inter-specific model, the reported non-significant statistic refers to the interaction of that effect with species.

Intercept-only models based on datasets containing only groups of interest were created to confirm if the group had retained the association, i.e. if they were choosing significantly differently from random. Post-hoc comparisons between pairs of trials in the 8-day memory assay and between species per trial in the 3-day memory assay inter-specific model were conducted using the estimated marginal means from the R package *emmeans* v1.8.8 [73], and were corrected for multiple comparisons using the Tukey correction.

