## Supplementary Material for "Cognitive robustness in a new insect model of extended longevity"

#### Table of Contents

1. Supplementary Results ... 2
   No evidence of age-related decrease in starvation resistance in either species ... 2
2. Supplementary Figures ... 4
3. Supplementary Tables ... 6
   3.1 Summary of age-related memory impairment (AMI) studies in insects ... 6
   3.2 Sample size breakdown (species, age, trial) for cognitive experiments ... 13
      8-day long-term memory ... 13
      3-day long-term memory ... 13

### 1. Supplementary Results

***No evidence of age-related decrease in starvation resistance in either species***

The log-rank test showed that survival was lower in butterflies from the 2022 field season than the in butterflies reared in 2019, with median survival in 2019 of 62 days (maximum: 104 days) in *H. hecale* and 31 days (maximum: 54 days) in *D. iulia*, but a median survival in 2022 of 46 days (maximum: 80 days) in *H. hecale* and 21 days (maximum: 68 days) in *D. iulia* (Figure S1; *H. hecale*: $\chi^2_1 = 8.5$, $p = 0.003$, *D. iulia*: $\chi^2_1 = 30.9$, $p < 0.001$). However, sub-populations were subjected to the testing protocol at different ages, likely introducing confounding factors influencing survival due to the regular bouts of food deprivation. Additionally, butterflies surviving long enough to complete the experiment were either dissected upon completion (as in the 2019 cohort) or sometimes moved to unmonitored cages (as with *H. hecale* in the 2022 cohort). All such individuals were censored from this point, violating the assumption of non-informative censoring necessary for survival analysis. It is thus inappropriate to draw conclusions about inter-cohort differences in these groups, though they likely stem from differences in the condition of the wild-caught individuals from which stocks were established during each field season. The plots in Figure S1 are therefore presented for illustrative purposes, to justify the choice of age groups for each species and cohort, as it was clear during sampling that the reduced survival in the 2022 *D. iulia* cohort would require a reduction in the age of the oldest age group tested in this species (see Methods).

Considering the evidence for decreased starvation resistance with age in other insects [1, 2], we tested for a potential reduction in butterfly survival resulting from the 4-hour bout of food deprivation imposed during the cognitive assays. Kaplan-Meier survival analysis was used to generate estimates of survival probability the day before and the day of the naïve preference trial for each age group of each species. As this trial was the first time that individuals were subjected to starvation stress, a reduction in survival on this day could be indicative of vulnerability to starvation in that particular group. The reduction in survival was considered to be significantly different if the 95% confidence intervals for estimated survival between the two days did not overlap. There was no evidence for an increase in mortality during the naïve testing protocol in either species, neither in the younger age groups nor as the butterflies aged (Figure S2), therefore showing no evidence for decreased starvation resistance with age. This suggests that Heliconiini butterflies are generally robust to short bouts of food-deprivation, as individuals were only deprived of food during the mornings on testing days, with all butterflies provided food by 12:30pm at the latest.

#### 2. Supplementary Figures

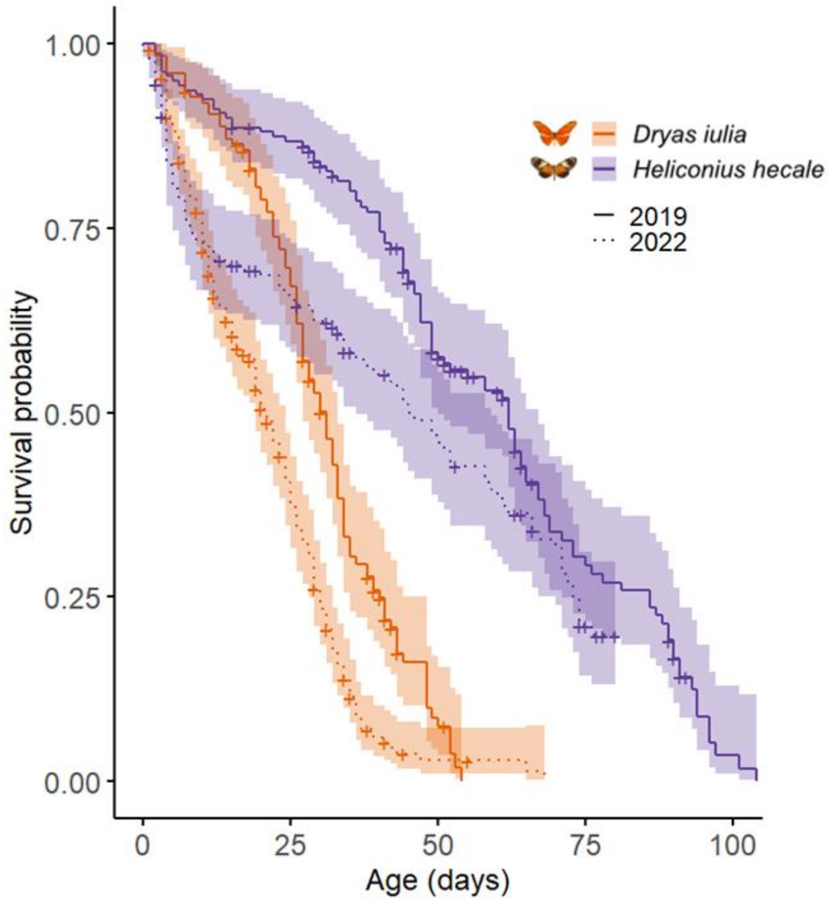


**Figure S1: Survival curves for *H. hecale* and *D. iulia* from the 2019 and 2022 cohorts.** Kaplan-Meier survival estimates and 95% confidence intervals for each species/cohort. “+” indicates a censored data point.

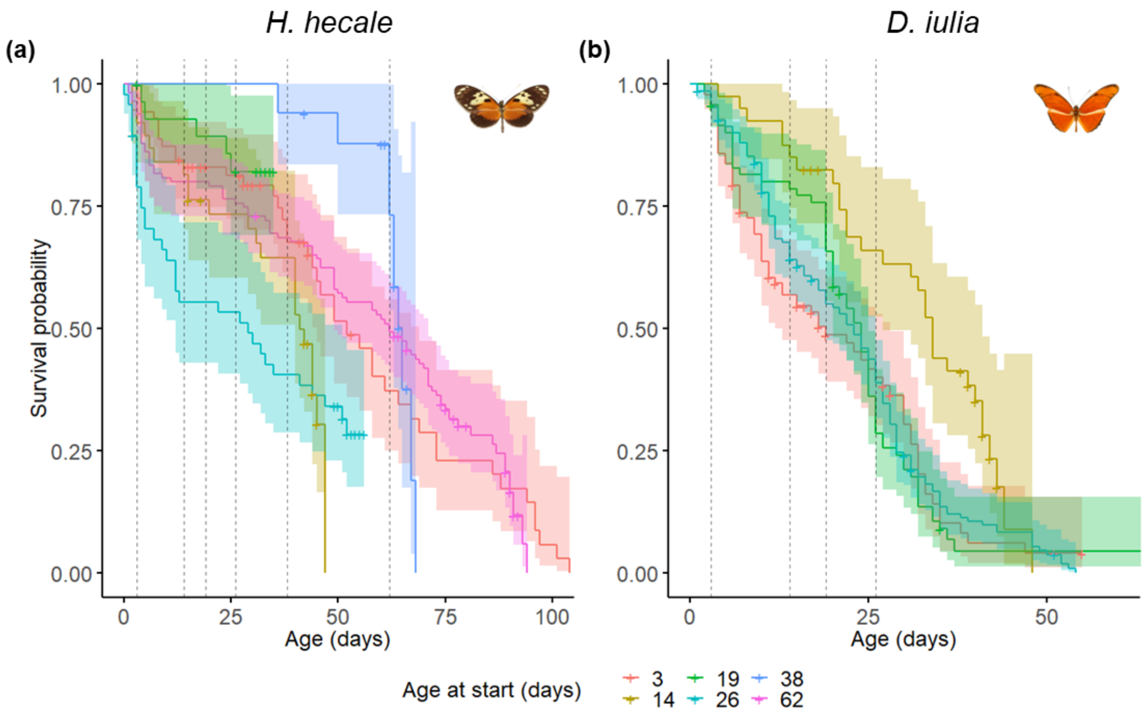


**Figure S2: Survival curves for *H. hecale* and *D. iulia* for each age group.** Kaplan-Meier survival estimates and 95% confidence intervals for (a) *H. hecale* and (b) *D. iulia* for each age group. Dotted vertical lines delineate ages at which separate groups were entered into the testing protocol, corresponding to the age groups for each species (see Methods). Any potential effects of starvation would in theory manifest as a reduction in survival after being first entered into the protocol. "+" indicates a censored data point.

### 3. Supplementary Tables

#### *3.1 Summary of age-related memory impairment (AMI) studies in insects*

**Table S1:** Summary of findings of all available studies of age-related memory impairment in insects to date (to the best of our knowledge). Learning and memory phases are far better characterised in *Drosophila* than in any other insect model, with described memory phases including short-term memory (STM), intermediate-term memory (ITM), anaesthesia-resistant memory (ARM), and protein-synthesis-dependent long-term memory (PSD-LTM) [3]. STM and ITM are both non-consolidated forms of short-term memory, whereas ARM and PSD-LTM are both consolidated forms of long-term memory. ARM is induced in *Drosophila* following a massed training protocol (although appears to also be induced at least in some cases following a single training cycle, see [4], is protein-synthesis-independent (and as such is sometimes referred to as PSI-LTM) [5, 6], is resistant to cold-shock anaesthesia, and persists for up to 3 days. PSD-LTM is induced in *Drosophila* following a spaced training protocol, is protein-synthesis-dependent, and persists for at least up to 7 days. Species are presented in order of decreasing numbers of studies, and then alphabetically by their binomial names.

| Species | Memory phase | Modality | Age of onset | Notes | Reference |
|---|---|---|---|---|---|
| Fruit fly (*Drosophila melanogaster*) | Learning (0 hours after training) | Olfactory | 10 days | Conditioning involved a single training cycle. Effect size was small and does not increase upon further ageing; authors suggest this might not truly be age-dependent. | [4] |
| | | | 25 days, further decrease at 50 days | Conditioning involved a single training cycle. | [7] |
| | Working memory (4 seconds after association) | Visual | 4 weeks – impairment<br>6 weeks – complete loss | Assayed “working memory” – retention for 4 seconds immediately following association. | [8] |
| | Short-term memory (3-20 minutes after training) | Olfactory | 21 days, 50 days | Tested 5 minutes after training, which involved 1-3 training cycles. Age was not the primary focus of this study, which was instead designed to look at the impact of dietary restriction on learning and memory – but they also report that short-term memory performance declined in old flies. | [9] |

| | | | | | |
|---|---|---|---|---|---|
| | | | No impairment (tested at 37-39 days) | Tested 20 minutes after training, which involved both massed and spaced training cycles. Age is described as age post-egg-laying, which for Drosophila is within 1-2 days of eclosion. | [10] |
| | | | 10 days | Tested 3 minutes after training, which involved a single training cycle. | [11] |
| | Intermediate-term memory (ITM) (1-3 hours after training) (non-consolidated, anaesthesia-sensitive) | Olfactory | 20 days – steady decrease up to 50 days | Tested 1 hour after training, which involved a single training cycle. Referred to in this paper as “middle-term memory” (MTM). However, also reported no impairment in 7h memory in flies of up to 50 days old. | [4] |
| | | | 21 days, 50 days | Tested 1 hour after training, which involved 1-3 training cycles. Age was not the primary focus of this study, which was instead designed to look at the impact of dietary restriction on learning and memory – but they also report that intermediate-term memory performance declined in old flies. | [9] |
| | | | 15 days | Tested 1 hour after training, which involved a single training cycle. Declines seen as early as 10 days in flies with antioxidant genes knocked down. | [12] |
| | | | 30 days | Tested 1 hour after training, which involved a single training cycle. However, also reported no impairment in 3h memory in flies of up to 50 days old. | [11] |
| | | | 25 days, further decrease at 50 days | Conditioning involved a single training cycle. Impairment in memory retention shown when tested both 1h and 3h after training. | [7] |
| | Anaesthesia-resistant memory (ARM) (3-24 hours after training) | Olfactory | No impairment (tested at 20 days) | Tested 3 hours after training. Conditioning involved a single training cycle; however, confirmed that this was ARM using cold-shock anaesthesia. | [4] |

| | | | | | |
|---|---|---|---|---|---|
| | | | No impairment (tested at 37-39 days) | Tested 24 hours after training, which involved massed training cycles. Age is described as age post-egg-laying, which for Drosophila is within 1-2 days of eclosion. | [10] |
| | | | No impairment (tested up to 50 days) | Tested 24 hours after training, which involved massed training cycles. | [11] |
| | | | 25 days, further decrease at 50 days | Tested 3 hours after training. ARM induced by subjecting flies to massed training cycles and confirming that memory was resistant to cold-shock anaesthesia. | [7] |
| | | | No impairment (tested at 30 days) | Tested 24 hours after training, which involved massed training cycles. | [13] |
| | Protein-synthesis-dependent long-term memory (PSD-LTM) (>24h) | Olfactory | 37-39 days | Tested 24 hours after training, which involved spaced training cycles. Age is described as age post-egg-laying, which for Drosophila is within 1-2 days of eclosion. | [10] |
| | | | 20 days | Tested 24 hours after training, which involved spaced training cycles. | [11] |
| | | | 25 days; further, steeper decline at 50 days | Tested 24 hours after training, which involved spaced training cycles. | [7] |
| | | | 30 days | Tested 24 hours after training, which involved spaced training cycles. | [13] |
| Honey bee (*Apis mellifera*) – summer bees | Learning | Olfactory (conditioned proboscis extension reflex in response to an odour) | No impairment (tested from 5 days to end of lifetime) | | [14] |

| | | | | | |
|---|---|---|---|---|---|
| | | | No impairment related to age | Tested “older” v.s. “younger” forager bees, average ages 32 v.s. 24 days. Found that deficits in olfactory learning were linked to social role rather than chronological age; older foragers displayed age-related decline in olfactory learning, but not age-matched nurses. Deficits can be compensated for by reversion of social role. | [15] |
| | | | No impairment (tested up to 52 days old) | Found no evidence for an impairment in learning; Behrends *et al*., 2007 argue that they did not control for behavioural function, e.g. if bees were workers/foragers etc. | [16] |
| | | | No convincing impairment related to age (technically age unknown, >15d foraging) | Presented evidence for an impairment in learning, but again seems more related to duration of foraging. Evidence for an actual age-related memory impairment is less convincing as for many individuals, actual age is unknown (“old” individuals in this study have simply spent more time foraging). | [17] |
| | | | No impairment related to age (chronological age not mentioned) | Confirmed deficit in olfactory learning in “old” foragers (longer foraging duration), but not age-matched nurses, and that these deficits could be reversed when foragers reverted to nurses. Confirmation that deficits appear related to social role rather than age. | [18] |
| | | | No impairment related to age | Demonstrated evidence of impairments in tactile learning, following same groups as [15], and again confirmed that deficits were linked to social role rather than chronological age, with deficits in foragers with long foraging durations but not age-matched nurses. | [19] |

| | | | | | |
|---|---|---|---|---|---|
| | | Tactile (conditioned proboscis extension reflex in response to a tactile pattern) | No impairment | Tested 72 hours after training. Found that long-term memory actually increased with age in longer-term foragers. | [19] |
| | Long-term memory (48-72 hours after training) | Tactile (conditioned proboscis extension reflex in response to a tactile pattern) | No impairment (technically age unknown, >15d foraging) | Tested 48 hours after training. Suggest disparity between their results and those in Drosophila may be due to differing experimental paradigms (here appetitive rather than aversive, timelines are different). | [17] |
| | | Olfactory (conditioned proboscis extension reflex in response to an odour) | No convincing impairment related to age (technically age unknown, >15d foraging) | Presented evidence for an impairment in spatial memory extinction, but again, this is unconvincing due to the correlation of these deficits with social role / foraging duration, and the unknown ages of the subjects. | [17] |
| | Extinction learning | Spatial | No impairment (technically age unknown, >15d foraging) | | [17] |
| | | Olfactory (extinction of conditioned proboscis extension reflex in response to an odour) | No impairment (tested at 6 months) | Argue in this paper that limited evidence for an impact of age on learning and memory suggests that the decisive factor in the manifestation of senescence in honey bees is the function of the individual in the hive (i.e. reduced activity state in the hive or active foraging outside the hive). | [20] |
| Honey bee (*Apis mellifera*) – winter bees | Learning | Tactile (conditioned proboscis extension reflex in response to a tactile pattern) | No impairment (tested at 6 months) | | [20] |

| | | | | | |
|---|---|---|---|---|---|
| | | Olfactory (conditioned proboscis extension reflex in response to an odour) | No impairment (tested at 6 months) | | [20] |
| | Short-term memory (5 minutes after training) | Tactile (conditioned proboscis extension reflex in response to a tactile pattern) | No impairment (tested at 6 months) | | [20] |
| | | Olfactory (conditioned proboscis extension reflex in response to an odour) | No impairment at 6 months | Nonsignificant trend that older winter bees performed worse than younger foragers, but not nurses, both 24 and 48 hours after training . | [20] |
| | Long-term memory (24-48 hours after training) | Tactile (conditioned proboscis extension reflex in response to a tactile pattern) | 160-180 days | Older winter bees performed worse than younger foragers/nurses 48 hours, but not 24 hours after training | [20] |
| | | Olfactory (conditioned proboscis extension reflex in response to an odour) | | | [20] |
| Two-spotted cricket (*Gryllus bimaculatus*) | STM (30 minutes – 2 hours after training) | Olfactory | No impairment (tested up to 3 weeks after final molt) | Tested 30 minutes and 2 hours after training, which involved multiple-trial conditioning. No impact of age was seen on either anaesthesia-sensitive nor anaesthesia-resistant memory, as confirmed by $CO_2$ treatment (tested 10 and 20 minutes after conditioning). | [21] |

| | | | | | |
|---|---|---|---|---|---|
| | LTM (4-24 hours after training) | Olfactory | 3 weeks after final molt | Tested 4 hours and 24 hours after training, which involved multiple-trial conditioning. | [21] |
| American cockroach (*Periplaneta americana*) | Learning | Visual (completion of a maze task) | 30 weeks; declined further at 50 weeks | Not classical conditioning; tested for ability to complete a maze task conducted daily over a 14-day period. | [22] |

##### *3.2 Sample size breakdown (species, age, trial) for cognitive experiments*

*8-day long-term memory*

**Table S2:** Sample sizes for analysis of 8-day long-term memory, listing the number of individuals for which data was collected for each age group in each trial. Sample size generally decreases as trials progress due to individuals dying before completing the experiment. Age (days) refers to the age at which individuals of that group were entered into the testing protocol. Trained = trained initial recall trial, LTM – long-term memory (suffixes 1-3 denote successive trials). As *D. iulia* are shorter lived than *H. hecale*, the oldest age point at which individuals of this species were entered into the testing protocol was 26 days.

| Species | Age (days) | *n* – Trained | *n* – LTM1 | *n* – LTM2 | *n* – LTM3 |
|---|---|---|---|---|---|
| *H. hecale* | 3 | 38 | 38 | 37 | 34 |
| | 14 | 24 | 22 | 21 | 18 |
| | 26 | 19 | 18 | 14 | 8 |
| | 38 | 15 | 13 | 11 | 4 |
| | 62 | 17 | 15 | 13 | 9 |
| *D. iulia* | 3 | 24 | 18 | 14 | 11 |
| | 14 | 24 | 15 | 11 | 8 |
| | 26 | 13 | 10 | 4 | 2 |

*3-day long-term memory*

**Table S3:** Sample sizes for analysis of 3-day long-term memory, listing the number of individuals for which data was collected for each age group in each trial. Sample size generally decreases as trials progress due to individuals dying before completing the experiment. Age (days) refers to the age at which individuals of that group were entered into the testing protocol. Trained = trained initial recall trial, LTM1 – long-term memory trial. The chosen age points were initially intended to correspond to those of the 8-day experiment, but overall survival was lower in this cohort, and so the oldest age point in *D. iulia*, and its corresponding age point in *H. hecale*, were reduced by a week to 19 days.

| Species | Age (days) | *n* – Trained | *n* – LTM1 |
|---|---|---|---|
| *H. hecale* | 3 | 23 | 19 |
| | 19 | 25 | 21 |
| | 62 | 21 | 13 |
| *D. iulia* | 3 | 23 | 16 |
| | 19 | 23 | 17 |
